# Crossing strategies of ecological barriers are affected by wing morphology and plumage colour in small migratory birds

**DOI:** 10.1101/2024.08.18.608464

**Authors:** Paul Dufour, Raphaël Nussbaumer, Martins Briedis, Pierrick Bocher, Greg Conway, Yannig Coulomb, Rose Delacroix, Thomas Dagonet, Christophe de Franceschi, Sophie de Grissac, Bastien Jeannin, Robin Monchatre, Fanny Rey, Stephan Tillo, Jocelyn Champagnon, Olivier Duriez, Frédéric Jiguet

## Abstract

The recent development of tracking technologies has revealed remarkable diel flight altitude changes over the Sahara Desert in small migratory bird species. However, the drivers and the traits that explain these crossing strategies remain poorly understood in part because so few species and barriers have been studied. Using a unique dataset from 67 recovered multi-sensor loggers, deployed across 17 species, we investigated when, where, and how birds cross two major marine barriers (Bay of Biscay, Mediterranean Sea) and a desert barrier (Sahara). Then, we relied on a comparative approach to examine the influence of wing morphology and plumage colour on these strategies. Our findings reveal important differences across barrier types. On average, birds fly at 1,600 m over the desert during nighttime, ascending to 2,800 m for prolonged daytime flights, while species crossing marine barriers fly significantly lower (750 m on average), in some cases flying just above the water surface during prolonged daytime flights. Wing morphology and plumage colour influence barrier-crossing strategies: flight altitude increases with wing area during both sea and desert crossings, and darker birds ascend to higher elevations during daytime Sahara crossings, likely to access cooler air and reduce solar heating. These findings refine hypotheses on barrier-crossing strategies and suggest broader ecological and evolutionary implications for migratory birds facing extreme environments.

## Introduction

Birds undertake some of the most spectacular annual migrations in the animal kingdom, with several billions of birds travelling twice a year between their breeding and wintering grounds [1]. In the Palearctic-African migratory corridor, it has been estimated that 2.1 billion songbirds and near-passerines migrate each season between Europe and sub-Saharan Africa [2], where they face several major ecological barriers. Ecological barriers are geographical features, such as deserts, mountains or seas, that can impact birds’ migration because they provide little food resources, nowhere to land or harsh climatic conditions [3] (see also [4]). Between Western Europe and sub-Saharan Africa, migratory birds can face marine barriers and the Sahara Desert which locally extends over 2000 kilometres. In some cases, birds can avoid and bypass these barriers, but under certain circumstances, they either choose or are forced to cross them [5].

How small migratory birds cross such large ecological barriers has long remained a mystery [6]. However, technological advances in recent decades, especially the miniaturization of tracking devices, have significantly expanded our understanding of their behaviours. Radar studies first provided evidence that small migratory birds mostly migrate at night with some birds regularly extending their flight into the day when crossing large barriers such as the Sahara Desert [7–10]. Light patterns recorded by light-level geolocators then confirmed that this strategy is probably common among small migratory species [11]. Finally, the use of multi-sensor loggers, including temperature, accelerometer and pressure sensors, has recently enabled accurate measurements of both the duration and altitude of these flights [12–14]. They notably revealed extremely high-altitude flights and diel cycles of flight dynamics during the Sahara crossing for different species (in *Acrocephalus arundinaceus* and *Upupa epops* [15,16]) and low daylight sea-crossing flights in *Caprimulgus europaeus* [17].

Several hypotheses have been proposed to explain these crossing strategies and mostly concerned desert crossings. They are related to factors such as predation, vision range, solar radiation, diel variation in ambient temperature, and wind support [16,18–20]. Regarding solar radiation, Sjöberg et al. [21] tested the hypothesis that birds fly at higher altitudes during daytime to mitigate the effect of extra heating from solar radiation. Using temperature data recorded by the loggers, the authors found that birds flying at the same altitude were warmer during the day that at night, which suggests that climbing during the day to colder altitudes might counterbalance the heating from solar radiation [19,21]. Few hypotheses have so far been proposed regarding marine crossings, partly due to the difficulty of tracking altitudinal movements and identifying such events in small species ([17], see also [22,23]).

Other factors, including wing morphology and plumage colour, may influence barrier-crossing behaviour, but they remain so far largely untested. Previous studies suggested that species with longer wings are more likely to undertake prolonged flights over the Sahara [12]. Longer and larger wings can indeed respectively enhance energy efficiency and lift [24–27], meaning some wing morphologies may not generate sufficient lift to reach higher elevations, where the air is thinner, when crossing barriers [28–30]. However, whether species with larger or more elongated wings tend to cross barriers at higher altitudes or complete longer, uninterrupted desert or marine crossings remain unclear. Regarding plumage colour, Delhey et al. [31] found that migratory birds generally have lighter plumage. Because darker birds absorb more solar radiation and heat up faster than lighter ones, authors suggested that migrants could have evolved reflective plumage to prevent overheating [32]. It remains to be tested whether darker species fly at higher altitudes, particularly during daytime, to mitigate excess heat from solar radiation. Testing these hypotheses have been challenging due to difficulties in tracking small species and pinpointing their exact locations during migration (see [33,34] for uncertainties of geo-positioning with light-level geolocators). While multi-sensor loggers have improved the geo-positioning [35], they still require recapturing individuals after their migration to retrieve data and few species have so far been equipped.

In this study, we equipped more than 300 individuals with multi-sensor loggers and analysed sea and desert crossings of 67 retrieved individuals of 17 small migratory bird species, that migrate between western Europe and their sub-Saharan wintering grounds. Our aim was to study how small migratory birds cross two major types of ecological barrier: marine and desert areas. First, we investigated whether barrier-crossing strategies differed among species, hypothesizing that morphological and plumage characteristics influence how individuals navigate ecological barriers. Specifically, we expected species with longer and larger wings to fly at higher altitudes and undertake longer flights [28,29], particularly over the Sahara, where the risk of overheating from solar radiation is higher. We also predicted that darker birds would ascend to cooler altitudes, especially during daytime crossings, to reduce heat absorption [21,31]. To test these predictions, we used a recently developed geo-positioning method that integrates activity, light, and pressure data [35,36], allowing us to accurately estimate where, when, and how each individual crossed the different ecological barriers. We then examined whether interspecific differences in flight altitude across the two barrier types were associated with morphometric traits and plumage colour.

## Materials and methods

### Geolocators and species data

For this study, 318 multi-sensor loggers were fitted on 17 species including 14 passerines and three non-passerines (see the list in Table 1) in different locations of France and the United-Kingdom (Figure S1) between 2021 and 2023. We used two types of loggers: 63 GDL3-PAM manufactured by the Swiss Ornithological Institute (1.2 g with harness) and 255 CARP30Z11-7-DIP manufactured by Migrate Technology (0.6 g with harness). The weight of the species tagged varied between 12 g for the smallest (*Acrocephalus schoenobaenus*) and 90 g for the heaviest (*Otus scops*) and the device always amounted to less than 5% of the body mass of the tagged birds. In the subsequent years, the same individuals have been recaptured to recover the tag. All birds in this study were breeding and experienced individuals that had completed at least one round-trip migration (see details in Supplementary Information).

**Table 1.** List of small migratory species tracked by multi-sensor loggers in this study. The number of loggers retrieved is indicated per species as well as the number of flights above the desert (Sahara Desert) and the marine (Mediterranean Sea and Bay of Biscay) barriers.

| Species (english) | Species (latin) | Loggers | Desert flights | Desert prolonged flights (>14h) | Marine flights |
| --- | --- | --- | --- | --- | --- |
| European Nightjar | *Caprimulgus europaeus* | 2 | 23 | 0 | 1 |
| Eurasian Scops-owl | *Otus scops* | 4 | 28 | 1 | 2 |
| Eurasian Hoopoe | *Upupa epops* | 4 | 23 | 2 | 5 |
| Woodchat Shrike | *Lanius senator* | 3 | 17 | 1 | 0 |
| Great-reed Warbler | *Acrocephalus arundinaceus* | 2 | 23 | 6 | 0 |
| Sedge Warbler | *Acrocephalus schoenobaenus* | 5 | 9 | 3 | 5 |
| Western Orphean Warbler | *Curruca hortensis* | 6 | 25 | 0 | 0 |
| Greater Whitethroat | *Curruca communis* | 1 | 2 | 1 | 1 |
| Spotted Flycatcher | *Muscicapa striata* | 5 | 15 | 7 | 0 |
| Common Redstart | *Phoenicurus phoenicurus* | 4 | 17 | 6 | 0 |
| Common Rock Thrush | *Monticola saxatilis* | 1 | 7 | 0 | 1 |
| Whinchat | *Saxicola rubetra* | 6 | 29 | 6 | 0 |
| Northern Wheatear | *Oenanthe oenanthe* | 7 | 36 | 3 | 9 |
| Common Nightingale | *Luscinia megarhynchos* | 3 | 12 | 2 | 1 |
| Western Yellow Wagtail | *Motacilla flava* | 7 | 16 | 13 | 9 |
| Tawny Pipit | *Anthus campestris* | 5 | 26 | 9 | 8 |
| Tree Pipit | *Anthus trivialis* | 2 | 4 | 4 | 0 |
| **Total** |  | 67 | 312 | 64 | 42 |

All loggers recorded ambient light intensity, activity data, atmospheric pressure, and temperature. Birds were captured and tagged under licenses delivered by the MNHN-CRBPO (reference program PP1190 Migralion and PP1245 Migratlane) and the British Trust for Ornithology, granted by the Special Marks Technical Panel (project code 2728).

### Description of barrier-crossing flights

To determine where, when and how each bird crossed the ecological barriers, we modelled the trajectory of each track following the approach presented in Nussbaumer et al. [36] and using the R package *GeoPressureR* (version 3.2). All steps needed to reproduce these estimates are described in the Supplementary Information. All data, code, and parameter values, including raw and labelled pressure, light, and activity data for each individual are available in [50]. This includes a *config.yml* file specifying individual settings such as deployment sites, crop dates, map extent and resolution, and the inclusion of wind or light data in movement model estimation.

We considered three barriers: the Sahara as a desert barrier (which can span up to 2,000 km) and the Mediterranean Sea and the Bay of Biscay as two marine barriers (respectively 500 – 800 km and 300 – 500 km). Using the most likely trajectory of each individual, we considered that a flight occurred over an ecological barrier if at least half of its duration happened within the polygon delimiting this barrier (see Figure S1). This threshold was chosen to ensure that most of the flight was meaningfully associated with the barrier environment, while accommodating potential uncertainty in trajectory estimation and barrier boundary delineation. Then, for each crossing flights, we extracted flight duration and median flight altitude, using barometric equation accounting for the pressure and temperature at ground level from the ERA 5 data at the most likely location. We also calculated the median flight altitude for the night (20:00 – 4:00 UTC) and the day periods only (8:00 – 16:00 UTC). We identified prolonged flight into daytime as flights lasting at least 14 hours (which generally corresponds to flights extending after 8:00 UTC). For prolonged flights, we also compared flight altitudes during night and day to test whether migrating birds tend to change altitudes between these periods. We used paired t-tests to assess whether the differences were statistically significant.

### Flight altitude differences among species

To quantify interspecific variation in wing morphology and plumage colour related to flight ability and altitude, we selected three explanatory variables: wing aspect ratio, hand-wing area, and plumage lightness. We obtained both wing aspect ratio and hand-wing area data from [28], and plumage lightness from [37], specifically focusing on the dorsal part of the bird, as it is the most exposed to solar radiation during migratory flights. When sex-specific plumage lightness values were reported in [37], we used the average value for the species to maintain consistency, especially since sex was not always determined for the individuals in our study. We verified the absence of collinearity between these variables and acknowledge that not directly recording wing morphology on captured birds can be seen as a limitation of our study.

After checking for phylogenetic relatedness in the data (see Supplementary Information), we fitted linear mixed-effects models using the package *lme4* [38] using all individual flight records and adding season (spring vs. autumn), species, individuals and taxonomy (Passeriformes vs. non-Passeriformes) as random effects. The last three random effects were structured hierarchically, with individuals nested within species and species nested within taxonomic groups. In cases where the model exhibited a singular fit, indicative of overparameterization and negligible variance in a random effect, we removed the random effect to improve model stability. For desert crossings, we tested separate models to assess how variables influenced median flight altitudes during the entire flight, night, and day periods, as well as flight duration, expecting different variables to affect flight behaviour depending on the period. For marine crossings, we examined only the relationship between the explanatory variables and the median flight altitude of the entire flight, since flight duration is largely determined by where the crossing occurs, and because few individuals flew during daytime.

## Results

A total of 67 multi-sensor loggers (6 GDL3-PAM and 59 CARP30Z11-7-DIP) were retrieved on 17 different species (Table 1). Analysis of the trajectories revealed 312 flights over the Sahara Desert and 42 flights over the Mediterranean Sea or the Bay of Biscay (see Table 1 and Figure S2).

We found significant differences of flight altitudes between species when crossing the Sahara (average of 1614 with a standard-deviation of 1048 meters above sea level; Figure 1A and Table S1). For example, *Lanius senator* (1033 ± 796 m asl, n = 17 flights) and *Phoenicurus phoenicurus* (1323 ± 1111 m asl, n = 17 flights) flew at lower altitudes than *Acrocephalus arundinaceus* (2466 ± 1420 m asl, n = 6 flights; Table S1). When flights extended into daytime, we found that 12 out of 14 species (85%) climbed to higher elevations compared to their night-time flight altitudes (paired t-test for all flights: t = 5.11, df = 76, p-value < 0.001; average of 2630 ± 1521 m asl; Figure 1A, 2). Among the 25 flights in which a clear change in altitude between night and day was visually detected, 68% of climbs began before the onset of civil twilight, and 36% occurred within the 20 minutes preceding civil twilight. Interestingly, three species, *Caprimulgus europeus, Curruca hortensis* and *Monticola saxatilis* did not exhibit any prolonged flights during daytime (see Figure 1C). Half of the flights over the Sahara lasted the duration of the night, i.e., between 10 and 14 hours (50%; Figure 1A). Flights lasting less than 10 hours were less common (30%), and about 20% of flights extended beyond the first night, lasting up to 45 hours. Very few birds stopped during the second day. When flights continued beyond one night and the following day, birds typically remained in flight until at least the morning of the second night.

**Figure 1.**
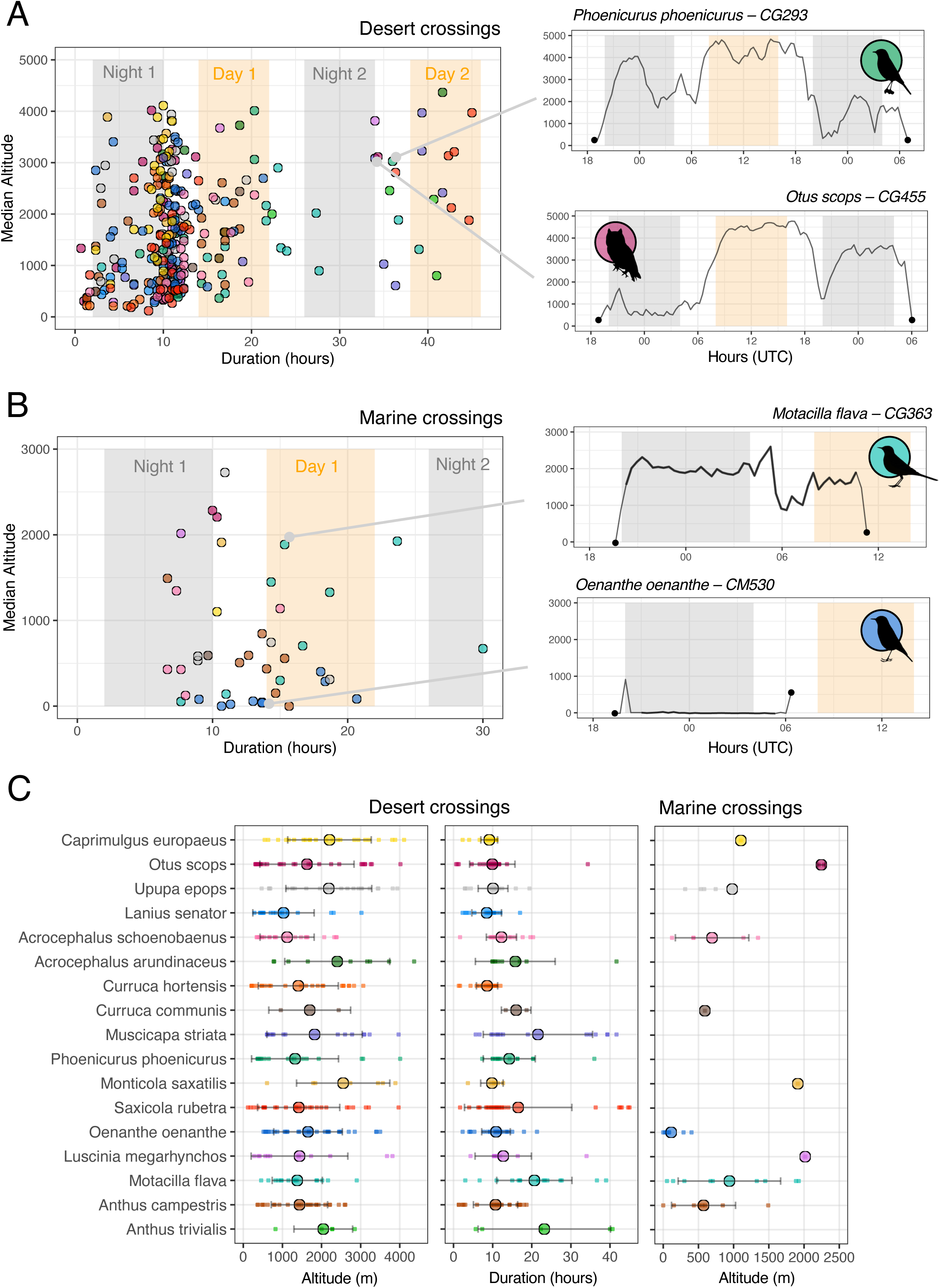
Variations in durations and altitudes of flights of small migratory species when crossing (A) the desert (Sahara Desert) and (B) the marine barriers (Mediterranean Sea and Bay of Biscay). Flight durations (in hours) are plotted against median altitudes (meters above sea level). Each dot represents a flight, coloured by species. Yellow and grey bars represent diurnal (8:00–16:00 UTC) and nocturnal (20:00–4:00 UTC) segments included in analyses. For marine crossings, the thicker lines in the two altitude profiles denote flights over the sea. In (C), average values per species of median altitudes and flight durations are presented for both desert and marine crossings. Flight durations for sea crossings are not plotted because they are largely determined by geographical distance rather than bird behaviour. Large dots represent species-averaged values, while small dots represent individual flights. Error bars represent standard deviation across individual flights. Silhouettes were downloaded from phylopic.org.

For sea crossings, flights generally lasted between 10 and 20 hours, depending on the sea extent to cross (Figure 1B). In addition, several individuals performed flights at very low altitudes, close to the sea surface (average of 774 ± 747 m asl). We found that 19 of the 42 marine crossing flights (45%) took place at a median altitude of less than 500 meters above sea level. In particular, *Oenanthe oenanthe* performed flights at an average altitude of 112 ± 137 m asl (n = 9 flights) and *Anthus campestris* at 572 ± 455 m asl (n = 8 flights). When flights extended into daytime, we found that 4 out of 5 species (80%) descended to lower elevations (paired t-test for all the flights: t = -3.55, df = 35, p-value = 0.001; Figure 2). Altitude profiles of all prolonged desert and sea crossings are shown in Figures S3 and S4, respectively.

**Figure 2.**
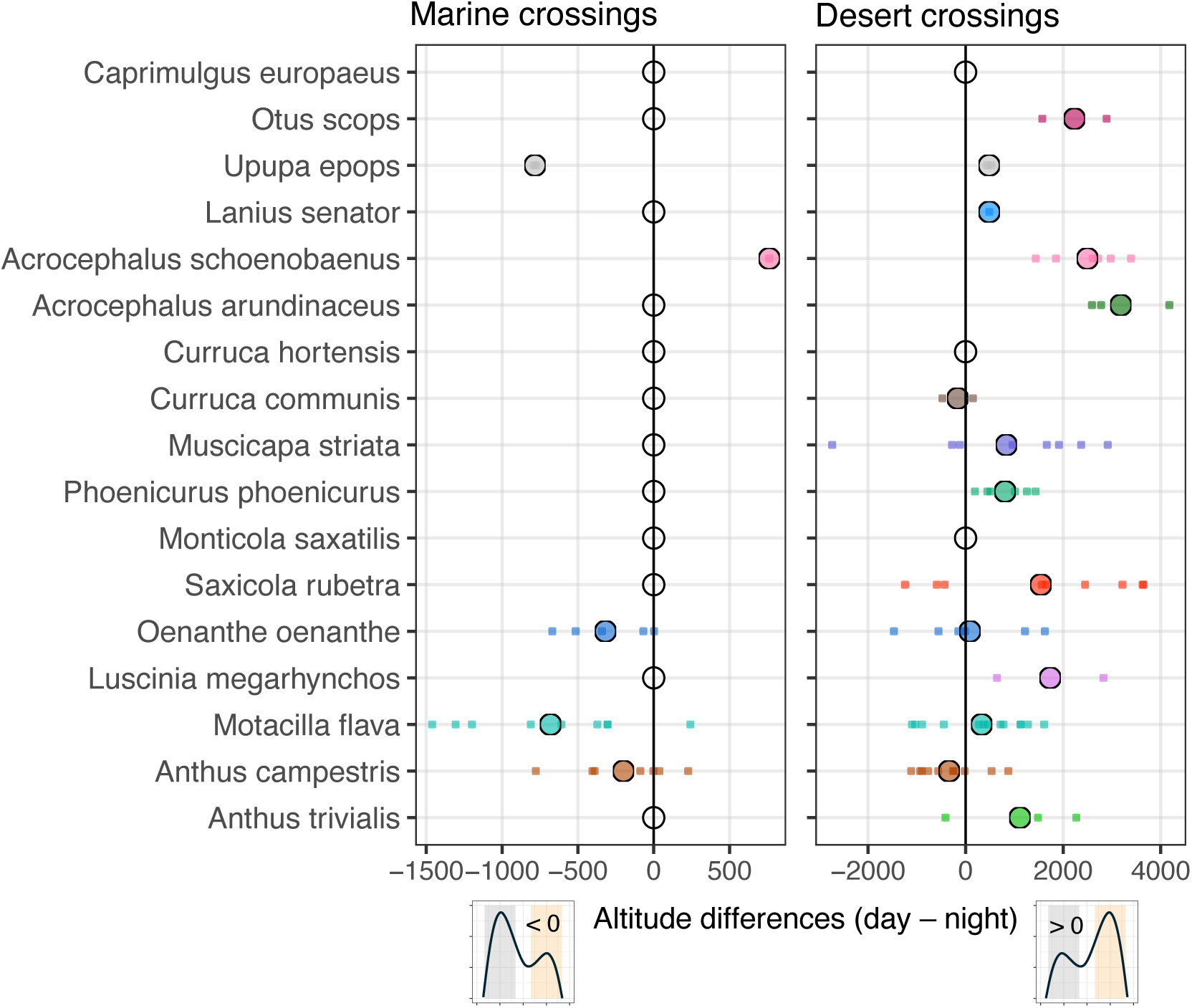
Differences in flight altitude between day and night of small migratory species when crossing marine and desert barriers. The x-axis indicates the altitude difference (day minus night), with positive values signifying higher flight altitudes during the day and negative values indicating higher altitudes at night. The coloured dots represent mean values. Empty (unfilled) dots denote species that did not engage in prolonged flight during daytime.

As we found no phylogenetic signal in the PGLS models (λ < 0.001, Table S2), we focused on linear mixed-effects models, which account for intra-specific variation by including multiple individuals per species and multiple flights per individual (Table 2). For desert crossings, hand wing area was positively associated with median flight altitude during both the entire flight and the night period (Table 2, Figure 3), while aspect ratio and plumage lightness were not significant. During the day period, median flight altitude showed a negative relationship with plumage lightness and a positive one with hand wing area; aspect ratio remained non-significant. Only hand wing area was significantly and negatively related to flight duration. Variance partitioning showed substantial contributions from individuals, species and taxonomy (nested together), and season across most models (Table 2). For marine crossings, individuals, species, and taxonomy again contributed substantially to variance, while season had minimal influence. Median flight altitude was significantly associated with all three traits: aspect ratio and hand wing area showed positive relationships, and plumage lightness a negative one (Table 2).

**Figure 3.**
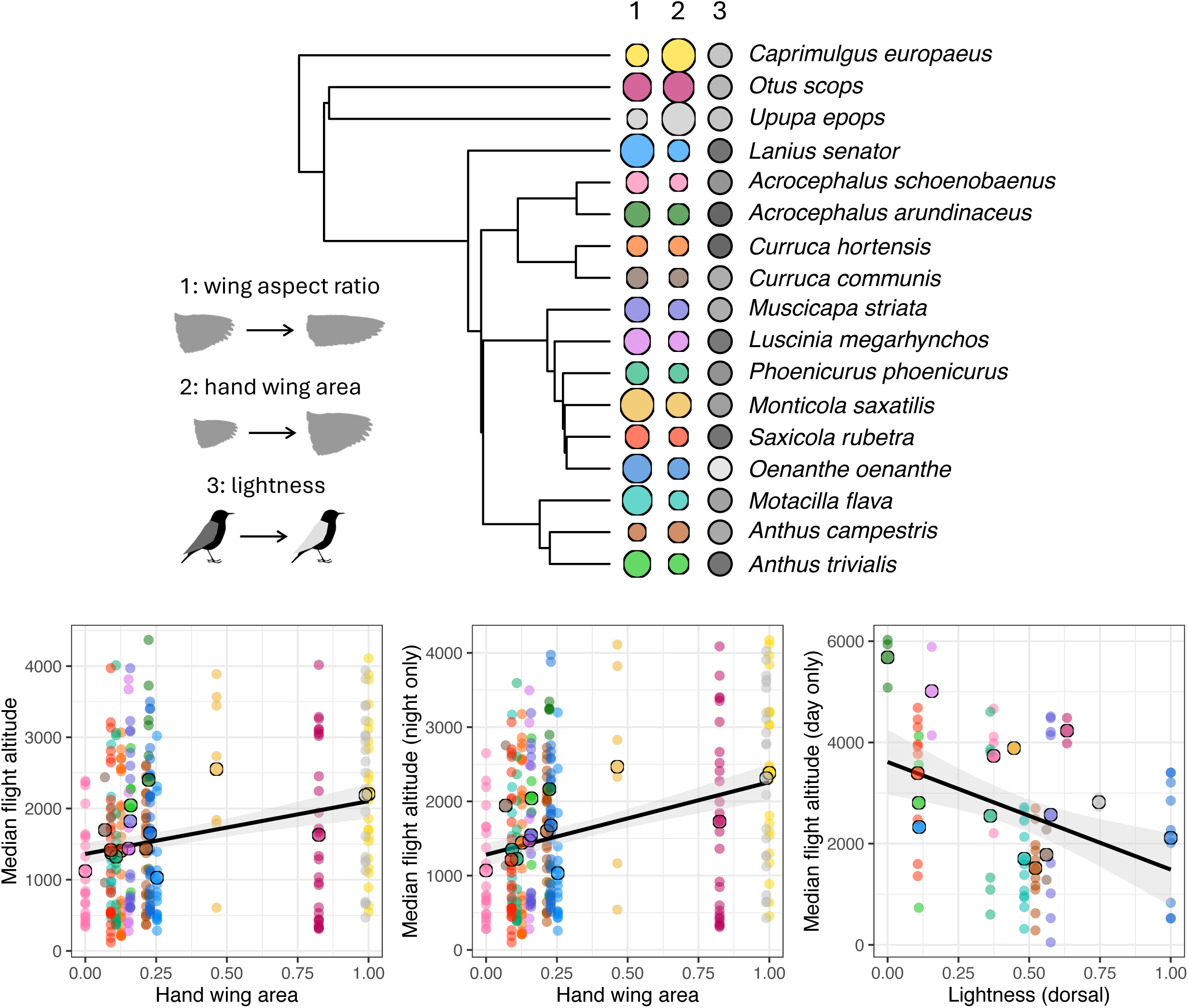
Relationship between wing morphology, plumage colour and flight altitude. The upper panel shows a maximum clade credibility tree of the studied species, with coloured circles whose size and shading represents variations of three morphological traits: (1) wing aspect ratio, (2) hand wing area, and (3) lightness of the dorsal part of the plumage. The lower panels display scatter plots linking the median flight altitude to hand wing area, the median flight altitude of the night period to hand wing area and the median flight altitude of the day period to the plumage lightness both during desert crossings (see Table 1 for other relationships). The black line represents the best-fit linear regression line and the shaded grey area around the black line represents the confidence interval for the regression line. Large dots represent species-averaged values, while small dots represent individual flights.

**Table 2.** Results of the mixed-effects models analysing the influence of wing morphology and plumage colour on flight behaviour across different barriers (desert and marine barriers). Fixed effects include aspect ratio, hand wing area, and lightness of the plumage, while random effects account for season, individual, species-level and taxonomic (nested together as ind:sp:taxo) variation. Estimates (Est.), standard errors (Std. Error), and p-values (p.value) are reported for fixed effects, along with variance components for random effects. Significant effects (p < 0.05) are highlighted.

| Barrier | Response | Fixed effects |  |  |  | Random effects |  |
| --- | --- | --- | --- | --- | --- | --- | --- |
|  |  | Variables | Est. | Std. Error | p.value | Variables | Est. |
| Desert | Median altitude | Intercept | 1334,4 | 488,8 | 0,192 | ind:sp:taxo | 104028 |
|  |  | Aspect ratio | 54,2 | 240,9 | 0,822 | Season | 432927 |
|  |  | **Hand wing area** | **658,0** | **229,1** | **0,006** | Residual | 736081 |
|  |  | Lightness | 118,1 | 236,6 | 0,619 |  |  |
|  | Median altitude - night | Intercept | 1282,8 | 460,9 | 0,186 | ind:sp:taxo | 72731 |
|  |  | Aspect ratio | -111,6 | 228,1 | 0,623 | Season | 384751 |
|  |  | **Hand wing area** | **825,8** | **214,4** | **<0,001** | Residual | 739189 |
|  |  | Lightness | 215,7 | 224,2 | 0,339 |  |  |
|  | Median altitude - day | Intercept | 3004,1 | 590,8 | 0,011 | ind:sp:taxo | 374163 |
|  |  | Aspect ratio | 748,2 | 640,7 | 0,251 | Season | 335567 |
|  |  | **Hand wing area** | **2041,2** | **880,6** | **0,023** | Residual | 141815 |
|  |  | **Lightness** | **-2438,4** | **651,1** | **<0,001** |  |  |
|  | Flight duration | Intercept | 14,0 | 2,1 | 0,004 | ind:sp:taxo | 17,0 |
|  |  | Aspect ratio | 2,8 | 2,5 | 0,267 | Season | 4,4 |
|  |  | **Hand wing area** | **-6,3** | **2,4** | **0,013** | Residual | 49,3 |
|  |  | Lightness | 0,2 | 2,4 | 0,939 |  |  |
| Marine | Median altitude | Intercept | 994,3 | 244,0 | <0,001 | ind:sp:taxo | 131969 |
|  |  | **Aspect ratio** | **843,2** | **341,9** | **0,021** | Residual | 194347 |
|  |  | **Hand wing area** | **1431,8** | **334,6** | **<0,001** |  |  |
|  |  | **Lightness** | **-1851,8** | **389,8** | **<0,001** |  |  |

## Discussion

### High altitude flights and overheating avoidance over desert barriers

Our results show that crossing the Sahara varies greatly from one species to another and can vary within species. Firstly, we confirmed that some species often extend flights during the day or over multiple days, while others stop after the first night [11,12,16,39]. For example, *Curruca hortensis* never extends nocturnal flights into the day and likely stops in acacia-filled wadis during spring migration across the western Sahara [40–43]. Species like *Oenanthe oenanthe* and *Anthus campestris* also make short flights and may be more flexible in habitat use in the Sahara. In contrast, frequent day-flyers like *Muscicapa striata* may have greater endurance or stricter stopover habitat needs, making desert crossings more challenging. The negative relationship between hand-wing area and flight duration was unexpected, suggesting that species with smaller wings tend to fly longer. This contrasts with our hypothesis that elongated wings favour prolonged flight. One possible explanation is that environmental conditions during migration may outweigh the effect of wing shape [44–46]. Additionally, smaller-winged species might use more energy-efficient flight styles, such as flap-bounding or low-speed cruising, that extend flight duration.

Regarding flight altitude, we found important differences among species. Several species reached unexpected high altitudes, such as *Otus scops*, *Acrocephalus schoenobaenus* or *Saxicola rubetra*, which all flew higher than 4,000 m during daytime. Among species that extend flights into the day, most climb higher than the previous night, but some, like *Oenanthe oenanthe* and *Anthus campestris*, maintain or even descend to lower altitudes. Our analyses also demonstrate that wing morphology and plumage colour influence barrier-crossing strategies over the desert. Indeed, we found that hand wing area was positively correlated with flight altitudes, which corroborates that idea that larger winged birds can generate more lift with less effort, enabling them to sustain flight at higher altitudes where the air is thinner [28,47]. Additionally, we found that the lightness of the plumage was negatively correlated with flight altitudes considering the daytime period only. These results confirm the prediction of Delhey et al. [31] that because darker birds absorb more solar radiation than lighter ones, they would tend to fly at higher altitudes to reach cooler conditions and mitigate excess heat from solar radiation. These findings are consistent with the hypothesis proposed by Schmaljohann et al. [19] and tested using multi-sensor loggers by Sjöberg et al. [21], suggesting that birds fly at higher altitudes during the daytime to reduce the thermal load from solar radiation.

### Low altitude flights over marine areas

Among the 17 small, nocturnal migratory bird species, no single strategy emerged for crossing marine barriers, but we observed a frequent strategy among some species, such as *Anthus campestris* and *Oenanthe oenanthe*, to fly at very low altitudes, often just above the water’s surface, especially during daytime.

Our data show that several birds traveling at these low altitudes quickly reach a constant altitude, which they maintain for most of their crossing. Norevik et al. [17] found a similar behaviour of very low altitude flight during the daytime in *Caprimulgus europaeus* and suggested that this behaviour could substantially reduce flight costs. Near-water flights can save energy, particularly in situations of headwinds or crosswinds, where birds can mitigate these effects by flying in weaker winds close to the water surface [48] or by flying in ground effect, where the aerodynamic cost associated with induced drag is considerably reduced [49]. Interestingly, we observe this behaviour here in species whose flight styles are quite different from that of the nightjar (i.e., wagtails, pipits or hoopoe). A possible explanation is that this low sea-crossing strategy has been selected, independently of the aerodynamic flight styles, to save energy, thereby also preventing overheating when these migratory birds cross large marine areas.

Interestingly, we found that variables included as fixed effects significantly predicted the median flight altitude during marine crossings. Both aspect ratio and hand-wing area had a positive effect, while plumage lightness had a negative effect: a pattern similar to that observed during desert crossings, despite most marine flights occurring at night (with only some individuals continuing into daylight hours). We believe these results should be interpreted with caution given the limited sample size (42 flights), and that additional data are needed to better understand the factors driving altitude variation during marine crossings. While we hypothesize that our findings (e.g., low-altitude flight over marine areas) may hold in other geographic contexts, this remains to be tested. Tracking birds crossing the Gulf of Mexico, for instance, would offer a valuable comparison because migratory flights in that region are typically longer and may involve different wind conditions and energetic demands than those over the Mediterranean Sea or the Bay of Biscay [5].

### Limits and conclusions

Our study highlights substantial inter- and intra-specific variation in flight strategies across ecological barriers, with flight altitude influenced by wing morphology and plumage colour. However, some limitations must be acknowledged. We were unable to directly test other important factors known to shape migration strategies, such as wind conditions and sex. Additionally, sample sizes, especially for marine crossings, were limited. The low-altitude flights observed over the sea, particularly during the day, are of particular interest, but understanding their drivers will require further data. Finally, while the use of multi-sensor geolocators greatly improves trajectory reconstruction, we acknowledge that some uncertainty remains. However, we believe its impact on our results is limited, as barrier crossings were classified based on the crossing of broad polygons rather than precise locations. Despite these limitations, our findings refine current hypotheses on barrier-crossing behaviour and underscore the combined role of environmental and morphological drivers. Future studies incorporating broader taxonomic coverage, environmental data, and physiological measurements will be key to deepening our understanding of avian strategies in extreme environments.

## Acknowledgements

Barbara Helm, Pablo Capilla-Lasheras, Felix Liechti and Steffen Hahn for discussions at different stages of the project development and the writing of the paper. Mathieu Gravey for his help developing the GeoPressureR package for the analysis of this study. Thibaut Lacombe for his help to deploy and retrieve the loggers. The reviewers for their valuable comments and suggestions on earlier versions of this manuscript. This study is part of the Migralion and Migratlane projects, which are founded by the Office Français de la Biodiversité (OFB).

## Supporting Information

### Supporting Text

#### Materials and methods

##### Geolocators and species data

For this study, 318 multi-sensor loggers were fitted on 17 species including 14 passerines and three non-passerines (see the list in Table 1) in different locations of southern and western France and the United-Kingdom (only for *Anthus trivialis*) between 2021 and 2023. The list of all locations can be found in Supplementary Data and in Figure S1.

We used two types of loggers: 63 GDL3-PAM manufactured by the Swiss Ornithological Institute (1.2 g with harness) and 255 CARP30Z11-7-DIP manufactured by Migrate Technology (0.6 g with harness). The GDL3-PAM loggers recorded pressure data at 5-minutes intervals and activity data every 30 minutes (Liechti *et al*. 2018). The CARP30Z11-7-DIP loggers recorded both pressure and activity data every 20 minutes. Most devices were mounted using a leg-loop harness made with a stretchy elastic thread with a diameter depending on the size of the species (0.7 to 1.0 mm). Note that we used a tressed UV-proof thread for *Otus scops*, *Lanius senator*, *Caprimulgus europaeus,* and *Upupa epops* to prevent the bird from damaging the harness with its beak and hence avoiding the bird to get entangled in it or to lose the logger.

All birds were individually marked with a metal ring (mandatory) and an engraved colour ring to facilitate their detection for recapture the following year. The vast majority of individuals in this study were captured using vertical mist nets (of different lengths and size of mesh) but note that some *Oenanthe oenanthe* were also caught with spring traps. In certain cases, playback devices were used to attract the birds. All captured birds were equipped, measured (mass, wing length) before being released after ca. 10 min of handling. Note that we did not include sex in our analyses because (1) not all individuals were sexed in the field, either because it was accidentally missed or due to difficulty in determining their sex, and (2) sex ratios were highly skewed across the studied species, with females being generally rare in the captures and, in some cases, only males being captured, likely due to their stronger responsiveness to playback.

##### Geolocators data analyses

To determine where, when and how each bird crossed the ecological barriers, we modelled the trajectory of each track following the approach presented in Nussbaumer et al. (2023a) and using the R package GeoPressureR (version 3.2). We followed all the steps presented in the online tutorial: https://raphaelnussbaumer.com/GeoPressureManual/

First, we separated stationary and flight periods by manually labelling the geolocator pressure and activity measurements. Stationary periods were characterized by a limited variation in consecutive pressure measurements indicating an absence of change in altitude associated with a limited variation in consecutive activity measurements. Migratory flights typically displayed a clear drop in atmospheric pressure, corresponding to altitude gain in-flight associated with consecutive high activity measurements (which forms a plateau of the same duration as the variation in pressure).

Second, we built separate probability maps based on atmospheric pressure and light intensity data (sunrise and sunset times) to estimate the position of each bird during each stationary period. Note that we did not use light intensity data (and only atmospheric pressure) for the nocturnal species *Otus scops* and *Caprimulgus europaeus* due to poor quality data (individuals spending most of daytime under cover). For the pressure-based maps, the time series of the geolocator pressure measurements during stationary periods were matched with the one-hour atmospheric pressure at surface level from ERA5 reanalysis dataset (spatial resolution: 0.5×0.5°) to produce a likelihood map of the geolocator’s position (following the method of Nussbaumer et al. 2023a). The resulting likelihood map included the information of both the temporal variation of pressure and the absolute values of pressure corresponding to the altitudinal range within each grid cell. For the light-based maps, we calculated likelihood estimates following Nussbaumer et al. (2023a). We used an “in-habitat” calibration from the equipment and retrieval periods (as described in Lisovski et al. 2012; Lisovski & Hahn 2012) fitting the distribution of zenith angle with a kernel density estimation. The likelihood maps of twilights belonging to the same stationary period were aggregated with a log-linear pooling.

Finally, we constructed the trajectories of each bird following a Hidden Markov Model presented in Nussbaumer et al. (2023b). The observation model consisted of the likelihood maps generated from pressure and light data. The movement model used the information of flight duration derived from the labelling and a parametric equation defined as the cubic root of the mechanical power required for the average airspeed computed for a transition, accounting for the different species size and shape. For all individuals, a low (flight) airspeed threshold of 20 km/h was used to account for potential short local or exploratory flights. Using this model, we generated (1) the marginal probability map of the position of each stationary period and (2) the most likely migration trajectory of the birds.

All the data and code to reproduce the trajectory estimation are available in Dufour (2025). The repository includes all the labelled pressure, activity, and, where applicable, light data files (light data are available only for certain species, as noted above). These labelled files are used either to distinguish between flight and stationary periods (activity data) or to identify pressure patterns in order to generate likelihood maps. In addition, the repository contains the raw (not labelled) data as well as a comprehensive .config file that lists, for each individual, all the parameters required to reproduce the analysis. This includes, for example: the location where the GLS device was deployed, the capture dates, the spatial extent of the map used to estimate the trajectory, and whether or not light data were used in addition to pressure data. We encourage readers to consult the accompanying tutorial for a detailed explanation of each parameter included in the configuration file.

Note that the folder on Zenodo is structured as a GeoPressureTemplate and contains all folders and files needed to re-produce trajectory outputs (as explained https://raphaelnussbaumer.com/GeoPressureManual/geopressuretemplate-intro.html).

##### Flight altitude differences among species

To test for phylogenetic relatedness, we used the package *phylolm* (Tung Ho & Ané 2014) to fit four phylogenetic generalized least square (PGLS) models while controlling for phylogenetic relatedness with Pagel’s λ lambda (Pagel 1999). We tested how the three explanatory variables were associated with the median flight altitudes for the entire flight, the night period only, and the day period only and the flight duration (all averaged per species) over the Sahara (see Supplementary Information). To do so, we constructed a phylogeny of the 17 species with a composite of bird phylogeny established by Prum et al. (2015) and a maximum clade credibility tree computed from 10,000 iterations of the Hackett backbone (Jetz *et al*. 2012), following the method of Cooney et al. (2017).

#### Results

A total of 67 multi-sensor loggers (6 GDL3-PAM and 61 CARP30Z11-7-DIP) were retrieved on 17 different species. Retrieving rates thus varied between 6.7% *Curruca communis* and 57.1% in *Otus scops* (24.1% on average). Specifically, we retrieved for example: 2 out of 26 tagged *Anthus trivialis* (7.2%; none of the loggers were recovered in France but 2 were retrieved in the United-Kingdom) ; 2 out of 21 tagged *Acrocephalus arundinaceus* (9.5%); 1 out of 3 *Monticola saxatilis* (33.3%); 5 out of 21 tagged *Anthus campestris* (23.8%); 3 out of 21 tagged *Lanius senator* (14.3%); 5 out of 10 tagged *Muscicapa striata* (50%); 5 out of 15 tagged *Acrocephalus schoenobaenus* (33.3%) and 4 out of 20 tagged *Phoenicurus phoenicurus* (20%). Returning rates, including individuals that returned but were not recaptured or lost their loggers, varied between 6.7% for *Curruca communis* and 57.1% for *Otus scops* (26.3% on average). Specifically, we had 3 *Anthus trivialis*, 1 *Lanius senator*, 2 *Motacilla flava*, and 1 *Upupa epops* that returned but were not recaptured, as well as 2 *Upupa epops* and 1 *Oenanthe oenanthe* that returned but without their logger. Finally, 2 *Motacilla flava* returned with their logger which stopped recording only after a few days, they were excluded from the study. A table detailing all the individuals equipped for this project, with the location of capture, the dates and biometry (at equipment and retrieval time, if retrieved) is available in Supplementary Information.

---

**Table S1.**
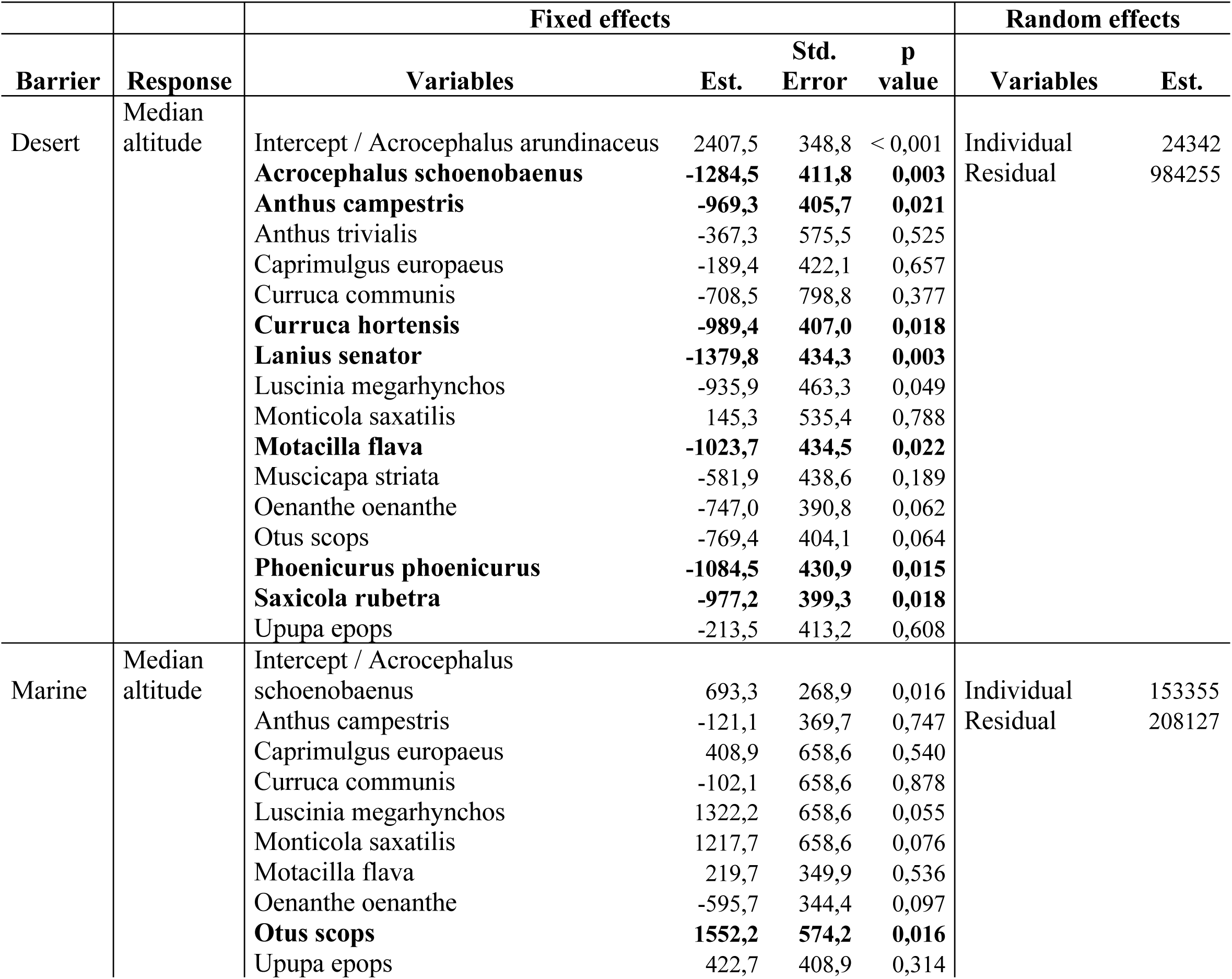
Results of the linear mixed-effects model testing for differences in median flight altitude between species for both desert and marine crossings. Species was included as a fixed effect, and individual ID as a random effect to account for repeated measures. Species with median flight altitudes significantly different from the intercept (p < 0.05) are represented in bold.

**Table S2.** Results of Phylogenetic Generalized Least Squares (PGLS) analyses modelling the effect of wing morphology and plumage colour on flight behaviour across desert crossings, using average values per species). With these models, we tested for the presence of phylogenetic signal before using linear mixed-effects models to include intra-specific variations (see methods and results).

| Barrier | Response | Variables | Est. | Std. Error | p.value | lambda |
| --- | --- | --- | --- | --- | --- | --- |
| Desert | Median altitude | Intercept | 1063,2 | 1223,2 | 0,409 | <0,001 |
|  |  | Aspect ratio | 70,04 | -6,3 | 0,728 |  |
|  |  | Hand wing area | 0,04 | 0,04 | 0,059 |  |
|  |  | Lightness | -3,3 | 0,2 | 0,784 |  |
|  | Median altitude - night | Intercept | 1223,2 | 1175,1 | 0,317 | <0,001 |
|  |  | Aspect ratio | -6,3 | 186,2 | 0,973 |  |
|  |  | Hand wing area | 0,1 | 0,01 | 0,024 |  |
|  |  | Lightness | 0,2 | 11,22 | 0,979 |  |
|  | Median altitude - day | Intercept | 3974,9 | 2631,3 | 0,159 | <0,001 |
|  |  | Aspect ratio | 300,9 | 447,5 | 0,515 |  |
|  |  | Hand wing area | 0,1 | 0,1 | 0,127 |  |
|  |  | Lightness | -6,3 | 20,2 | 0,002 |  |
|  | Flight duration | Intercept | 14,2 | 13,7 | 0,316 | <0,001 |
|  |  | Aspect ratio | 0,3 | 2,2 | 0,877 |  |
|  |  | Hand wing area | -0,1 | 0,1 | 0,099 |  |
|  |  | Lightness | 0,1 | 0,1 | 0,840 |  |

**Figure S1.**
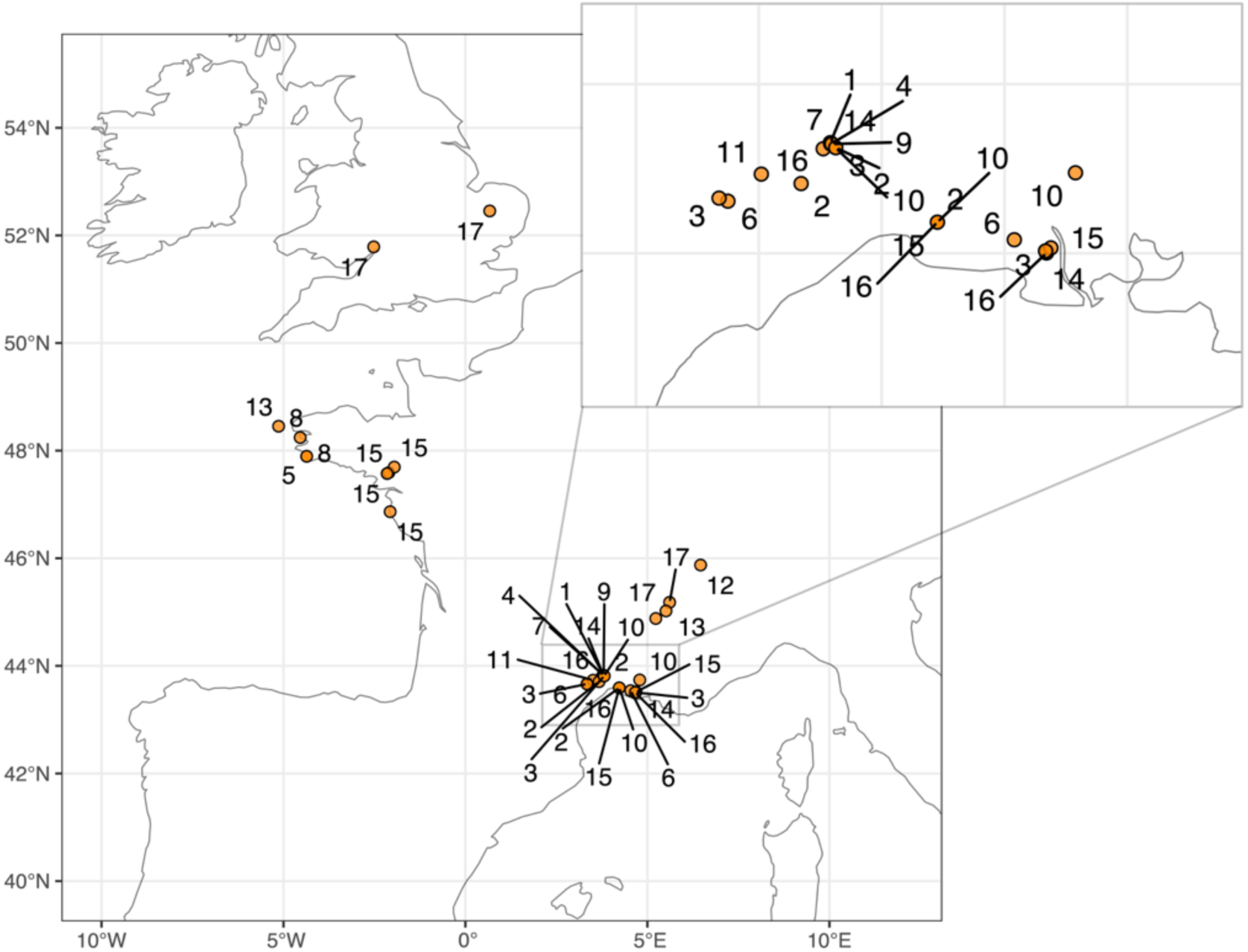
Capture sites of 17 bird species across Western Europe (France and the United Kingdom), with a zoomed subpanel showing the Montpellier/Camargue region in southern France. Species are numbered as follows: 1: Caprimulgus europaeus, 2: Otus scops, 3: Upupa epops, 4: Lanius senator, 5: Acrocephalus schoenobaenus, 6: Acrocephalus arundinaceus, 7: Curruca hortensis, 8: Curruca communis, 9: Muscicapa striata, 10: Phoenicurus phoenicurus, 11: Monticola saxatilis, 12: Saxicola rubetra, 13: Oenanthe oenanthe, 14: Luscinia megarhynchos, 15: Motacilla flava, 16: Anthus campestris, 17: Anthus trivialis.

**Figure S2.**
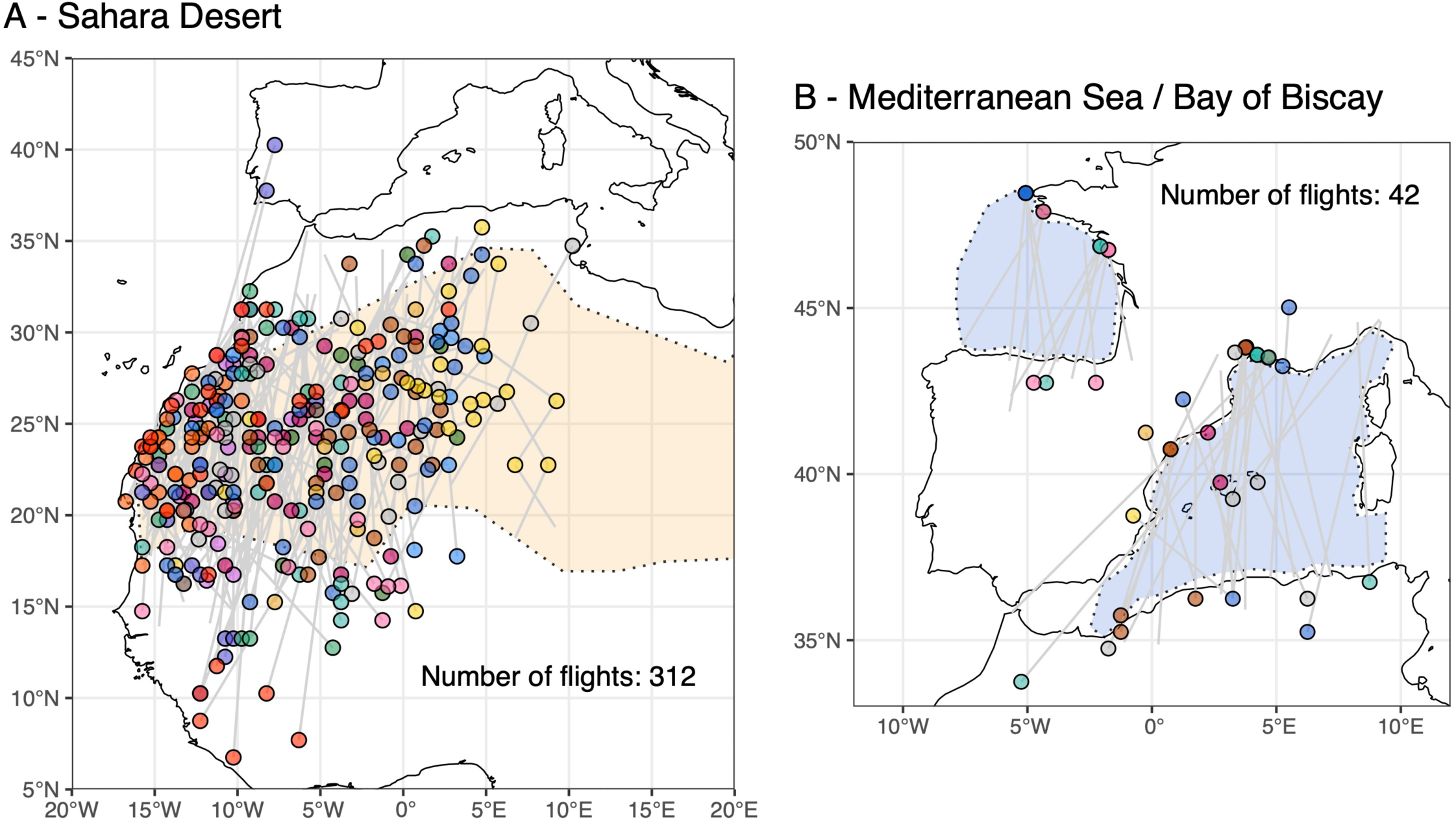
Locations of recorded flights over (A) the Sahara Desert and (B) the Mediterranean Sea and Bay of Biscay. Each flight appears as a grey line with a coloured dot marking the previous stop-over; colours denote species as in Figure 1. The orange polygon represents the Sahara Desert, while blue polygons indicate the Mediterranean Sea and Bay of Biscay. Flights are considered as crossing a barrier if at least half occurs within the defined polygon. For crossings of the Mediterranean Sea, we considered both direct crossings from the south of France or the north of the Iberian Peninsula to the African continent, as well as crossings that included a stopover in the Balearic Islands. For the Sahara Desert, we adopted the limits described in Brito *et al*. (2016).

**Figure S3.**
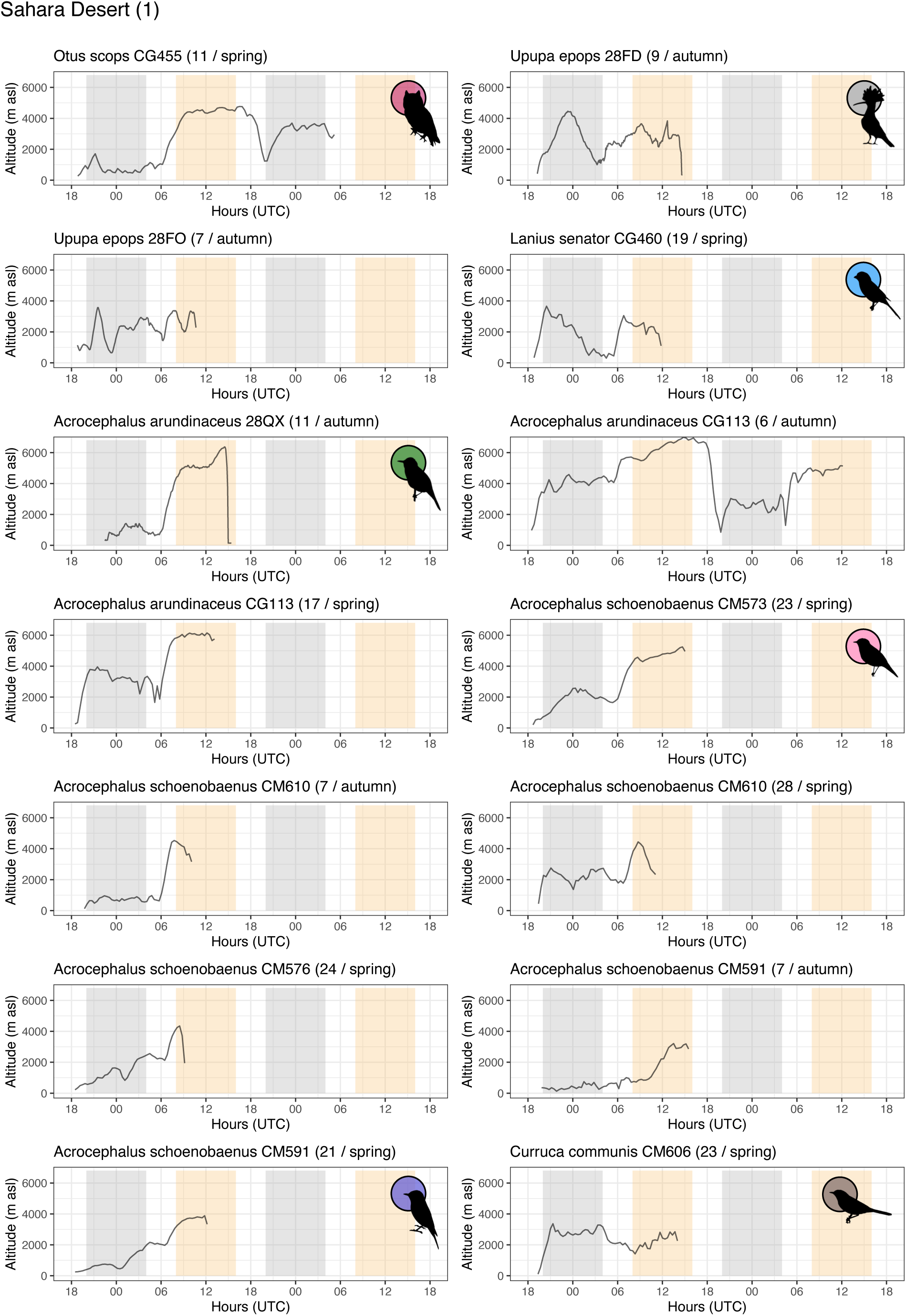
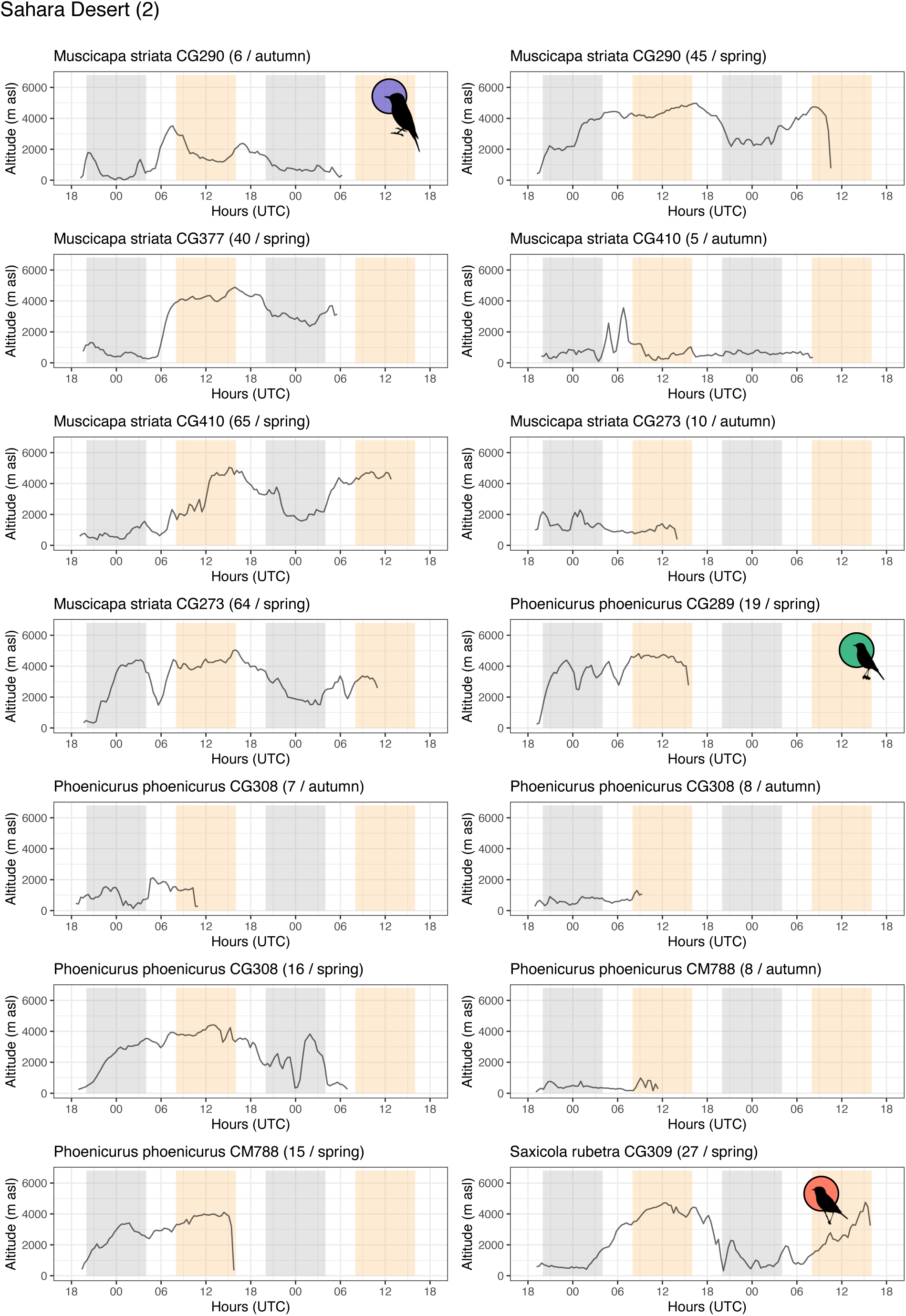
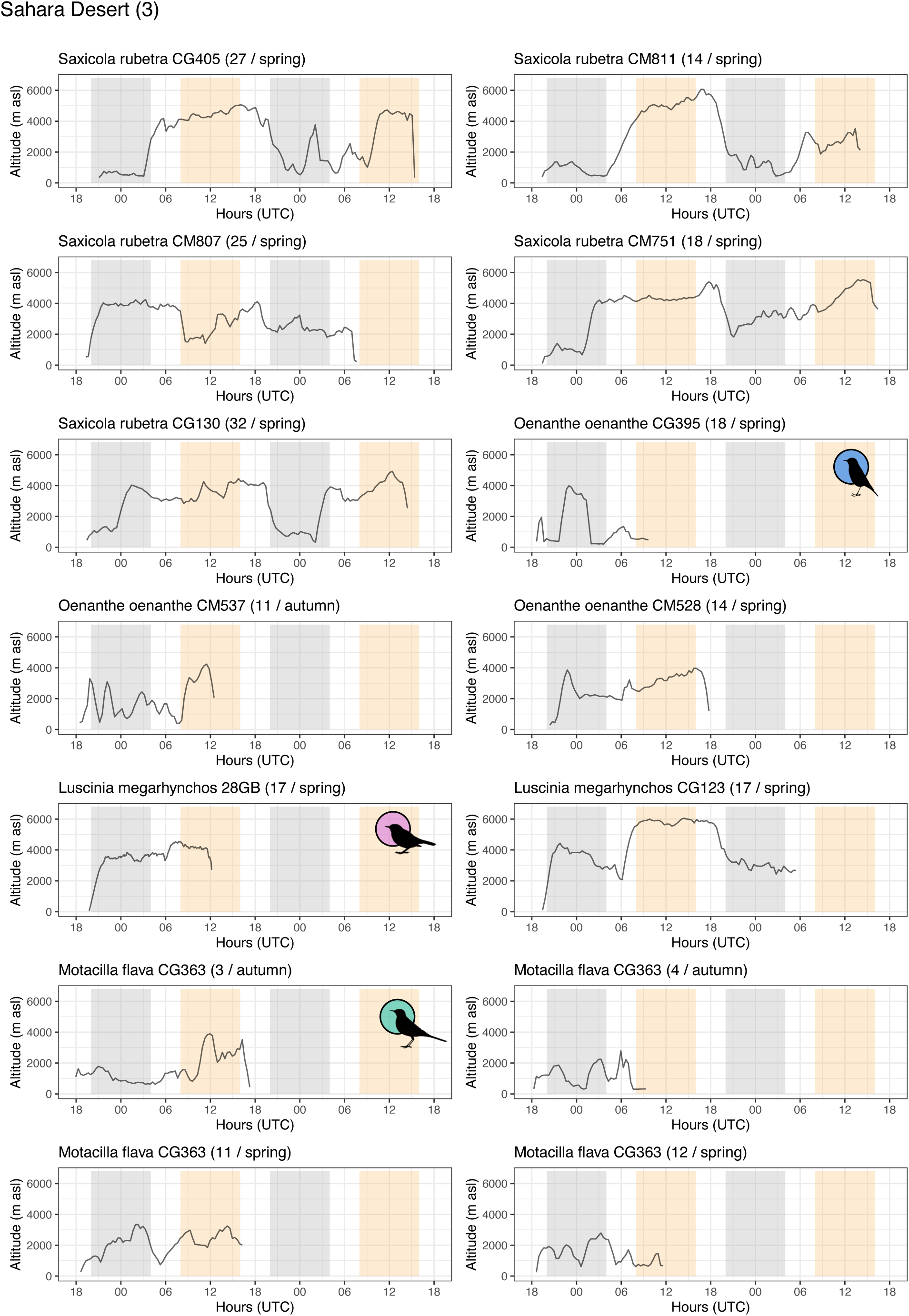
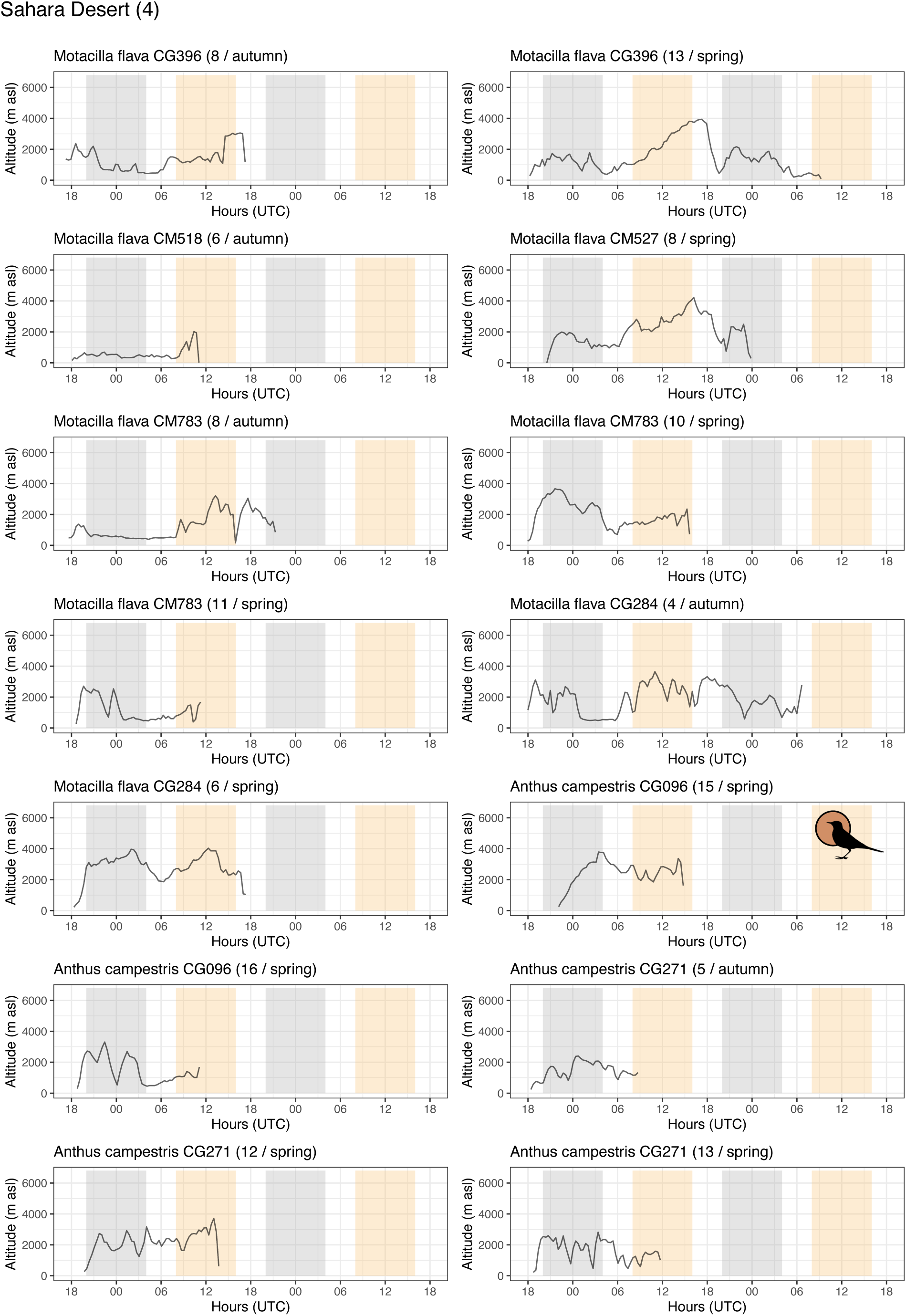
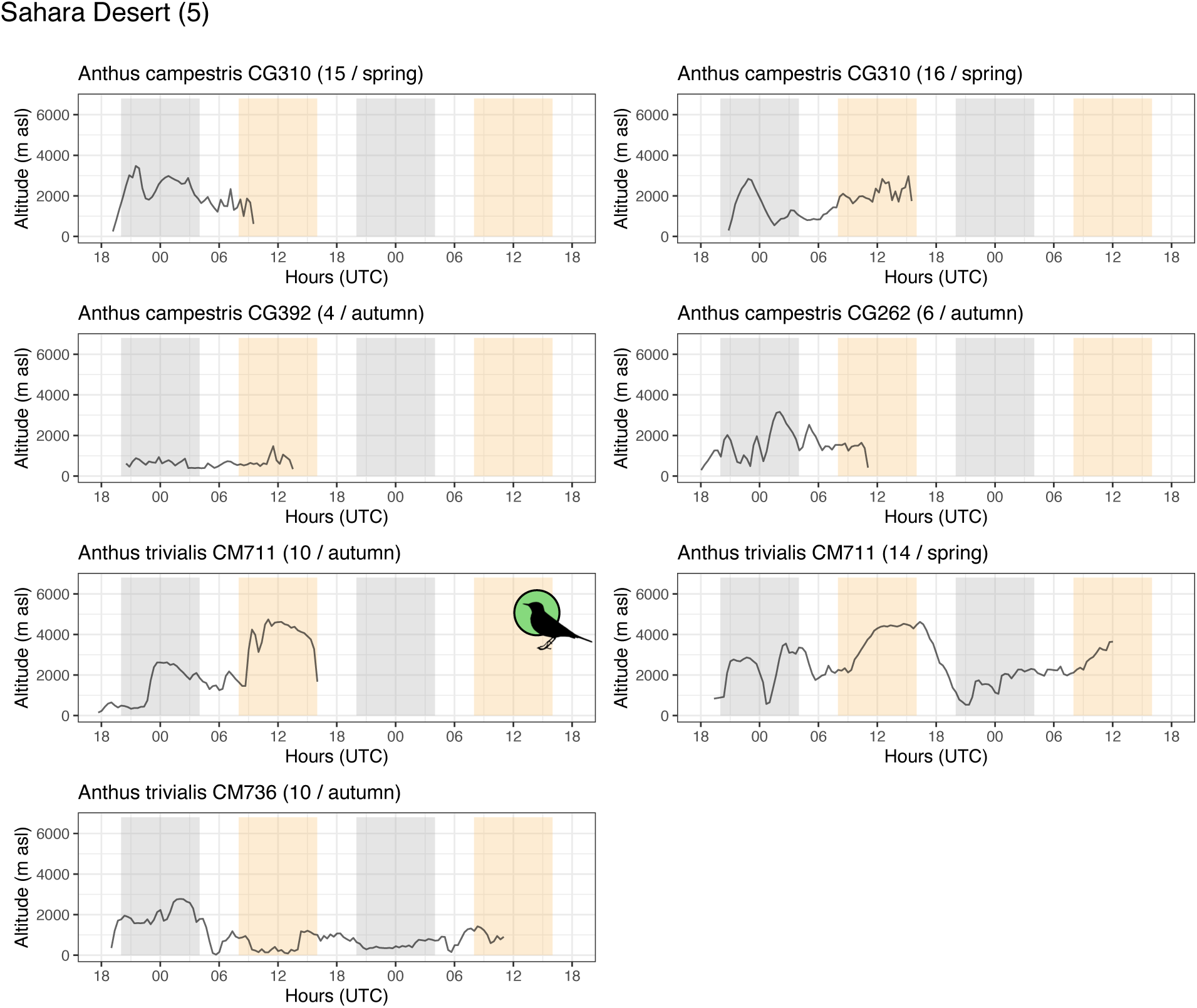
Variation in altitudes recorded among individuals of different species during prolonged flights (over 14 hours) across the Sahara Desert. On each panel, yellow and grey bars indicate parts of the flights that are included in the analyses as diurnal (8:00-16:00 UTC) or nocturnal (20:00-4:00 UTC). Silhouettes were downloaded from phylopic.org.

**Figure S4.**
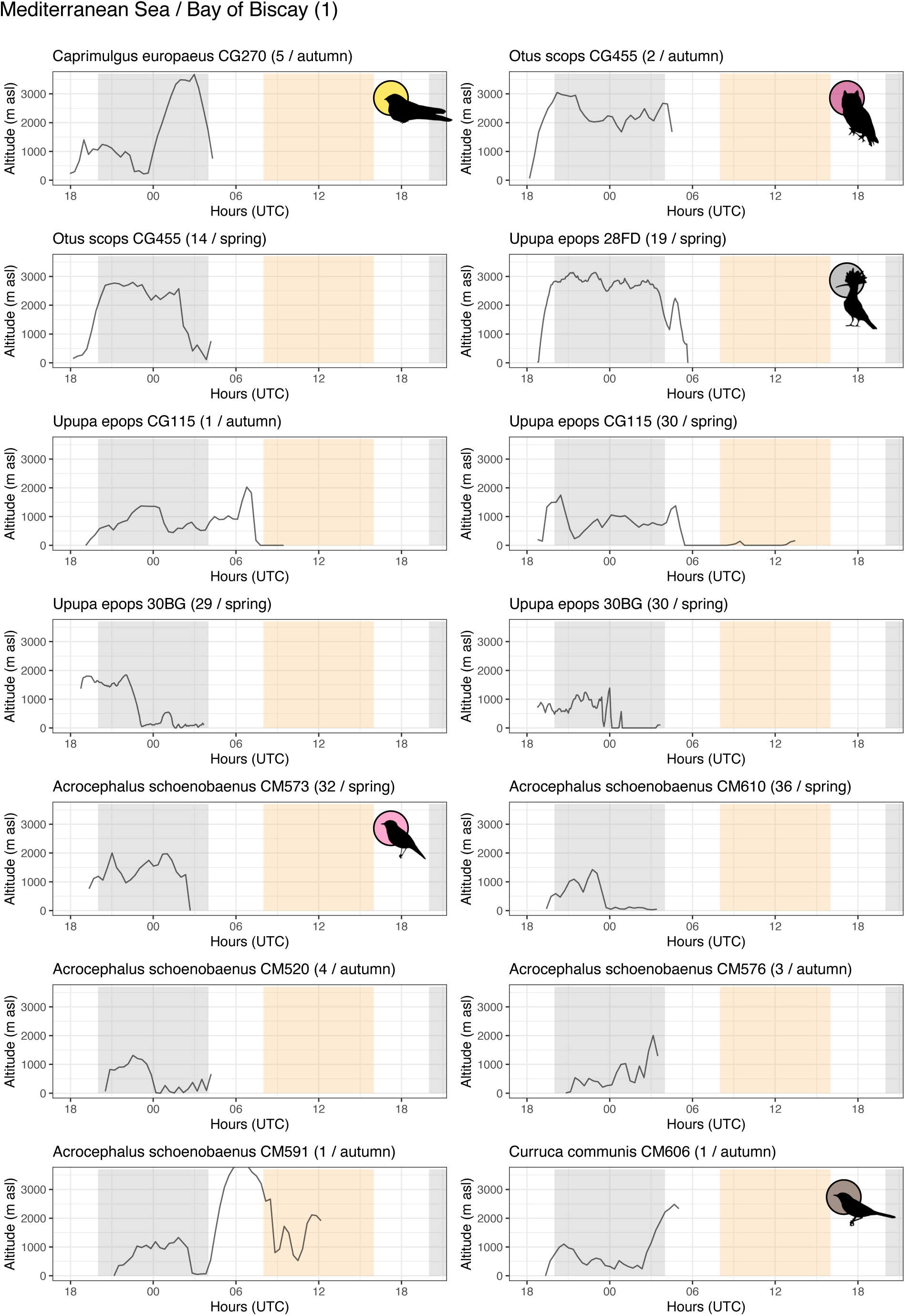
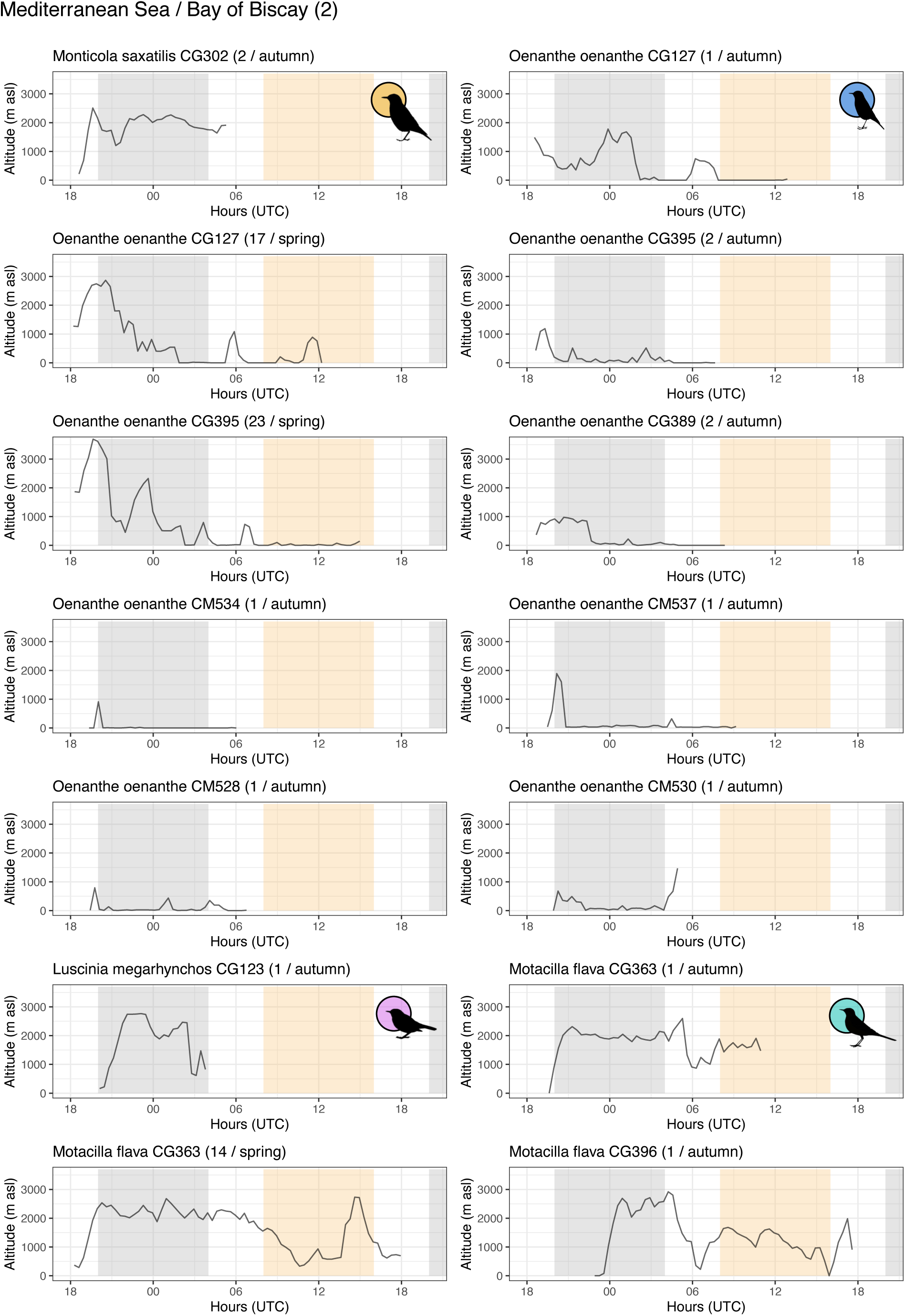
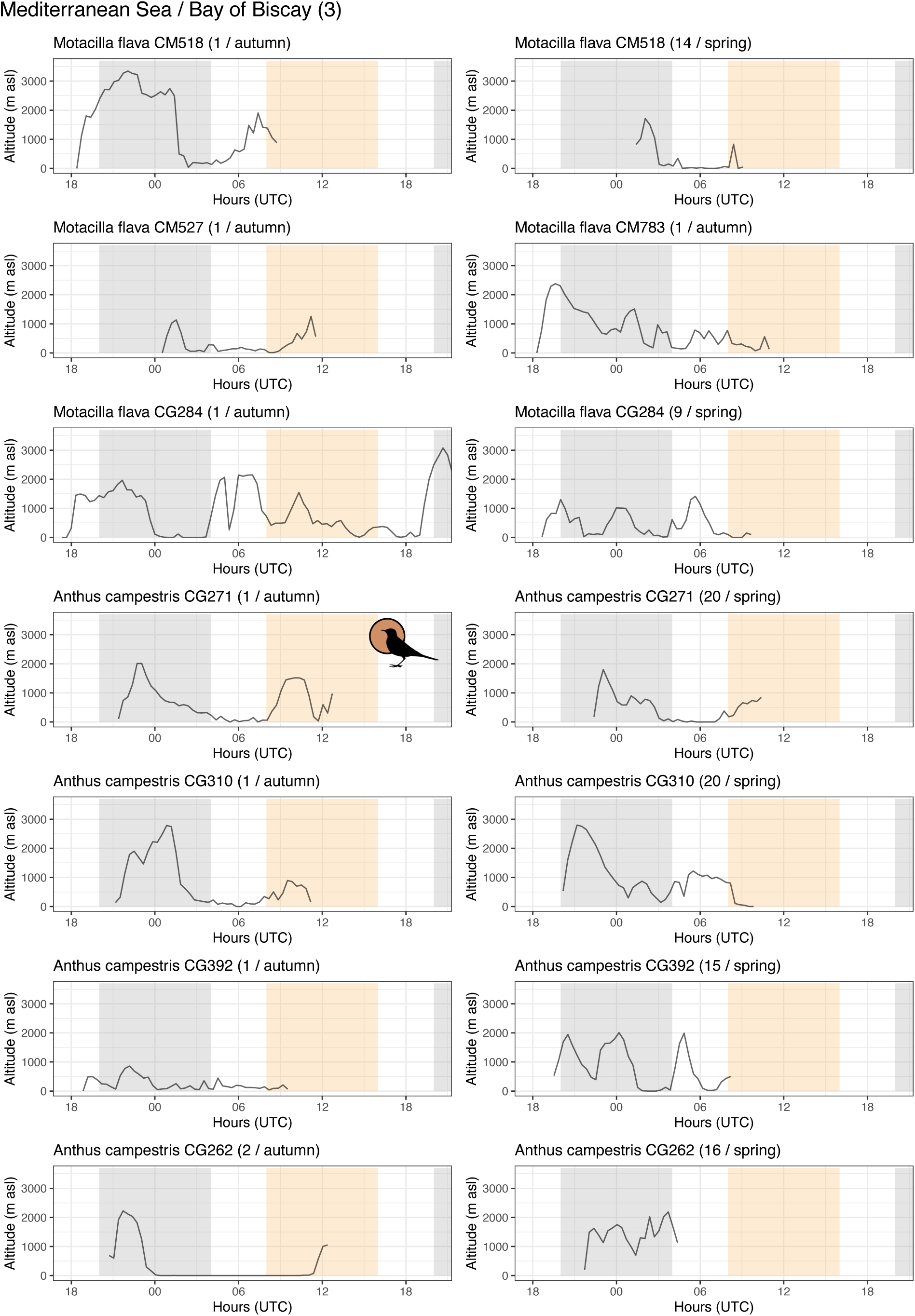
Variation in altitudes recorded among individuals of different species during sea crossing flights across the Mediterranean Sea and the Bay of Biscay. Flights across the Mediterranean Sea can be full crossing or interrupted crossing with a stop-over in the Balearic Islands. On each panel, yellow and grey bars indicate parts of the flights that are included in the analyses as diurnal (8:00-16:00 UTC) or nocturnal (20:00-4:00 UTC). Silhouettes were downloaded from phylopic.org.

